# Local adaptation shapes metabolic diversity in the global population of *Arabidopsis thaliana*

**DOI:** 10.1101/2021.09.13.460026

**Authors:** Rik Kooke, Willem Kruijer, Henriette D.L.M. van Eekelen, Frank F.M. Becker, Ron Wehrens, Robert D. Hall, Roland Mumm, Ric C.H. de Vos, Fred A. van Eeuwijk, Joost J.B. Keurentjes

## Abstract

The biosynthesis, structure and accumulation of secondary metabolites in plants are largely controlled by genetic factors, which can vary substantially among genotypes within a species. Here we studied a global population of *Arabidopsis thaliana* accessions for qualitative and quantitative variation in volatile and non-volatile secondary metabolites using essentially untargeted metabolomics. Genome-wide association (GWA) mapping revealed that metabolic variation mainly traces back to genetic variation in dedicated biosynthesis genes. Effect sizes of genetic variants, estimated by a Bayesian procedure, indicate that most of the genetic variation in the accumulation of secondary metabolites is explained by large-effect genes and defined by multiple polymorphisms. The various genetic variants resulted from independent mutation events and combined into distinctive haplotypes, which are representative for specific geographical regions. A strong relationship between the effect-size of regulatory loci, their allele frequencies and fixation index indicates that selection forces discriminate between haplotypes, resulting in different phytochemical profiles. Finally, we demonstrate that haplotype frequencies deviate from neutral theory predictions, suggesting that metabolic profiles are shaped by local adaptation and co-evolution of independent loci.

## Introduction

The success and evolution of life on earth relies almost completely on the ability of plants, as primary components of the food chain and net-producers of oxygen, to grow and flourish in a wide diversity of environments and conditions. For this, plants have adapted their morphology, developmental timing and metabolism to some of the most extreme settings (Cannell et al., 2020). Adaptation also includes the sometimes multi-trophic interactions with the biotic and abiotic environment. Most of the properties of plants, therefore, display an enormous variation in expression among and even within species. Although monogenic qualitative traits do occur (*e*.*g*., disease resistance or specialized organogenesis), the majority of traits are polygenic, or even omnigenic (Boyle et al., 2017, Chateigner et al., 2020). Such so-called quantitative traits have been shaped for millions of years through evolutionary processes, like mutation and recombination, drift, dispersal and natural selection, and their genetic architecture approaches an infinitesimal model, in which an infinite number of genes with infinitely small effects determines the resulting phenotype (Rockman, 2012, Olson-Manning et al., 2012).

*Arabidopsis thaliana* is no exception and is deservedly considered the reference plant species for studies on natural variation (Alonso-Blanco et al., 2009). *A. thaliana* diverged five to six million years ago from other species and despite relatively recent bottleneck events, due to glacial-interglacial climate changes, has adapted to a wide range of environmental settings spanning nearly all terrestrial habitats across the globe (Lyu, 2017, Shimizu and Purugganan, 2005). The species displays a wide diversity in the manifestation of traits and the abundant genetic resources available enabled the analysis of linkage between sequence diversity and natural variation in adaptive properties (Bergelson and Roux, 2010, Kover and Mott, 2012, Trontin et al., 2011). Being sessile organisms, unable to escape environmental threats, plants have evolved an enormous arsenal of phytochemical compounds, predominantly secondary metabolites, to combat risks and hazards (Fang et al., 2019). Natural variation in chemical profiles offers resilience to species and even increases evolvability in changing environments (Payne and Wagner, 2019, Segrè et al., 2002). Because plants need to be able to respond quickly to environmental fluctuations, a flexible and diverse chemical composition is advantageous and it is reasonable to assume that secondary metabolites are key targets upon which selection acts (Kooke and Keurentjes, 2012, Wu et al., 2018). Gaining insight into the genetic regulation of plant metabolites, being the last layer of the underlying molecular network, provides an alternative strategy to decipher the evolutionary forces that have shaped natural variation in quantitative traits (Chae et al., 2012, Kooke and Keurentjes, 2012).

Indeed, much of the metabolic disparity can be attributed to variation in genetic control and several mapping studies have associated local sequence diversity with metabolic traits and related higher order phenotypes (Chan et al., 2010, Fu et al., 2009, Joseph et al., 2015, Keurentjes et al., 2006, Keurentjes et al., 2008, Kliebenstein, 2009, Yu et al., 2020, Zhang et al., 2020). For instance, metabolic variation has been associated successfully with genomic diversity in genome wide association studies (GWAS) using linear mixed models (LMM), such as EMMAX (Fusari et al., 2017, Kerwin et al., 2015, Wu et al., 2016, Strauch et al., 2015, Li et al., 2020). Such approximate methods have demonstrated their effectiveness as a powerful tool to identify genetic associations, while considering relatedness among samples and accounting for population stratification and other confounding factors (Kang et al., 2010, Korte and Farlow, 2013, Lippert et al., 2011, Listgarten et al., 2010). However, GWA methods rely on frequentist statistics, which suffers from overfitting, bias in the estimation of effect sizes and low power of detecting rare alleles (Josephs et al., 2017, Zhou and Stephens, 2012). These drawbacks hinder the accurate estimation of the effect of genetic variation in an evolutionary context, which can be largely overcome by whole genome regression (WGR) methods, which include all genetic polymorphisms simultaneously into a single statistical model (de Los Campos et al., 2013). WGR provides accurate estimations of effect sizes of genetic polymorphisms, which is instrumental in the elucidation of the genetic architecture of traits and the selective forces acting on this.

Here, we aimed to determine the relative abundance of volatile and non-volatile secondary metabolites in a large global population of Arabidopsis accessions and provide accurate estimates of the effect of genetic polymorphisms on variation in these phytochemical profiles by applying a Bayesian WGR model (Moser et al., 2015). We further provide insight into the genetic architecture of metabolic traits by demonstrating that genes involved in metabolic biosynthesis pathways are the prime targets of evolutionary forces. Finally, we reveal that local adaptive processes and climatic variables shape global variation in plant secondary metabolism.

## Results

### Natural variation in plant secondary metabolism is driven by pathway diversity

To analyze the extent of natural variation in secondary plant metabolism, a global collection of 359 Arabidopsis accessions (Baxter et al., 2010, Horton et al., 2012, Li et al., 2010) was evaluated in duplicate for metabolic content in four-week-old rosettes of plants grown under standard conditions in short days. This population has previously been analyzed extensively for numerous morphological, metabolic and stress-related traits (Bac-Molenaar et al., 2015b, Davila Olivas et al., 2017, Fusari et al., 2017, Thoen et al., 2017, Wu et al., 2018, Kooke et al., 2016), although the systematic effect-size of genomic variation and genetic architecture was hardly established in these studies. Here, we investigated plant metabolic content in two independent replicate samples of each accession through both untargeted UPLC-Orbitrap-FTMS and SPME-GCMS based profiling, enabling the detection and relative quantification of non-volatile and volatile secondary metabolites, respectively (Salem et al., 2020, Kooke et al., 2019, Wehrens et al., 2016). A total number of, respectively, 567 and 603 non-volatile and volatile compounds were detected of which many could be putatively annotated using public and in-house databases (Perez de Souza et al., 2017, Tikunov et al., 2012, Kooke et al., 2019, Witjes et al., 2019). Widespread quantitative and qualitative variation in chemical profiles was observed among the genotypes of the population (Figure 1A; Supplemental Table 1). A large number of volatile and non-volatile metabolites were commonly detected in all accessions analyzed (32% and 17%, respectively), while 9% and 16% of the compounds were detected in less than 50 accessions, respectively (Figure 1C). For instance, each accession contained on average eight of the 94 rare non-volatile metabolites, but the accessions Zdr-6 and Mt-0 contained no less than 38 and 39 of these compounds, respectively. The lowest number (228) of non-volatile metabolites was detected in Cvi-0, although this accession contained the highest level of 3-butenyl glucosinolate. This pattern of variation in phytochemical profiles is consistent with earlier findings in a much smaller set of accessions (Keurentjes et al., 2006). Moreover, the variation in abundance of detected metabolites was substantial, with similar median coefficients of variation of 101% and 82% for volatile and non-volatile compounds, respectively. However, the broad-sense heritability (H^2^) of non-volatile metabolites was on average much higher than that of the volatile metabolites, possibly due to stronger stochasticity in the release of volatile compounds. Nonetheless, median heritabilities of 40% (volatiles) and 64% (non-volatiles) demonstrate that a large part of the variation in secondary metabolites is heritable (Figure 1B).

**Figure 1:**
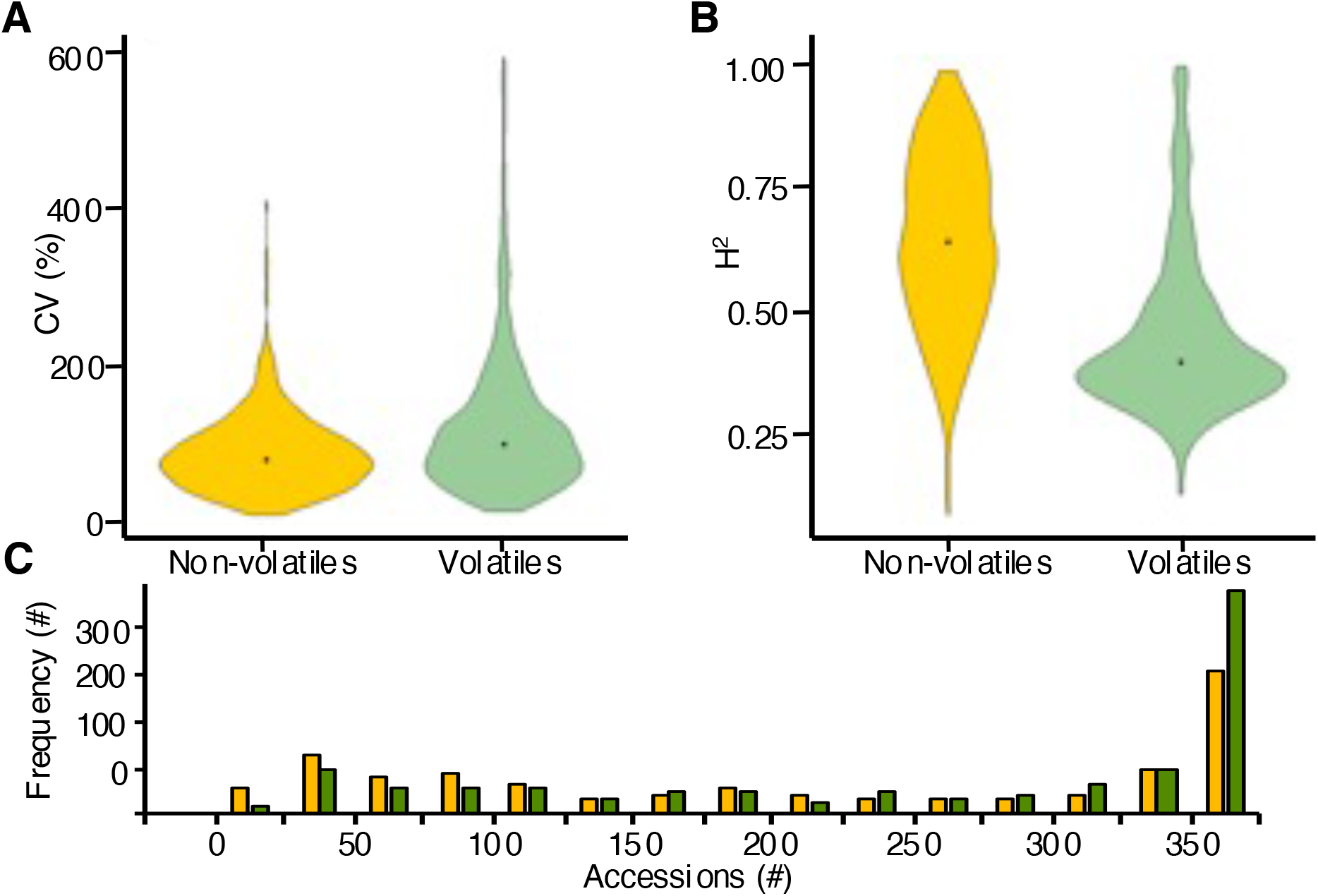
Genetic variation for metabolic accumulation in a global population of *Arabidopsis thaliana*. Distribution of the coefficient of variation (**A**) and the broad-sense heritability (**B**) of the relative abundance of volatile and non-volatile metabolites. (**C**) Frequency distribution of the number of accessions in which a specific metabolite was detected. Green and yellow items represent volatile and non-volatile metabolites, respectively.

With a view to identifying the causal genetic factors, we associated the observed variation in secondary metabolite profiles with ∼215K single nucleotide polymorphisms (SNPs) using a state-of-the-art Bayesian WGR method implemented in the Bayes-R software (hereafter referred to as WGR), which explicitly partitions the genetic variance into contributions of loci with large, intermediate or small effects (Moser et al., 2015). Since Bayesian WGR methods do not provide significance measures, a conventional GWAS using the EMMA-X approach (Kang et al., 2010) (hereafter referred to as GWA) was performed on the same data for reference purposes. Because WGR provides only SNP effect estimates, we set a conservative arbitrary effect size threshold of |*β*| > 0.01 for SNPs substantially contributing to explained variance, although the exact explained variance in metabolite accumulation depends also on the minor allele frequency (MAF) and total variance.

For roughly half of the 567 non-volatile metabolites (53%) at least a single SNP was detected that explained a substantial part of the variability (WGR: 300 metabolites (SNP effect size |*β*| > 0.01); GWA: 303 metabolites (SNP significance *P*_*BONF*_ < 0.05)). The proportion of the 603 volatile metabolites for which variation could be associated to one or more SNPs was substantially lower (WGR: 157 metabolites (26%, |*β*| > 0.01); GWA: 134 metabolites (22%, *P*_*BONF*_ < 0.05)) (Supplemental Table 2). The difference in mapping power between the two types of metabolites reflects their contrast in heritability, while the discrepancy between the WGR and GWA approach might be an effect of low minor allele frequencies. Rare large-effect loci are better detected using WGR while the power to detect true positives with LMM GWAS declines with decreasing MAFs. The latter would suggest that variation in volatile metabolites, in contrast to variation in non-volatile metabolites, is more frequently driven by rare alleles. The overlap in metabolites for which variation could be associated to genetic variation by both WGR and GWA was 83% for the non-volatiles and 57% for the volatiles, emphasizing the differences between metabolite types and mapping approach.

To assign genomic functions causal for the observed variation in metabolic profiles, candidate genes in linkage disequilibrium (LD) with the large-effect SNPs detected by WGR (|*β*| > 0.01) were identified. For many metabolites multiple QTLs were detected, often represented by series of large-effect SNPs in LD. In total, 1,097 unique candidate genes explaining variation in the 301 non-volatile metabolites were assigned, whereas 651 candidate genes were assigned explaining variation in the 154 volatile metabolites (Supplemental Table 2). Only 120 of the assigned genes explain variation in both non-volatiles and volatiles, illustrating that there is little overlap in the genetic regulation of volatile and non-volatile secondary metabolites.

At a number of loci, variation in several metabolites mapped to the same candidate gene. A QTL hotspot was defined as a gene locus substantially associated with variation in at least five metabolites by a minimum of two large-effect SNPs (|*β*| > 0.01). This criterion was met by nine loci after WGR analysis of non-volatile metabolites and three loci for volatile metabolites, with a small overlap of two hotspots detected with both platforms (Supplemental Table 2, Figure 2A). With the exception of *ACD6* and *GLABRA1*, which are involved in trade-offs between growth and defence (Fusari et al., 2017, Todesco et al., 2010) and trichome development (Wang et al., 2019, Herman and Marks, 1989, Marks and Feldmann, 1989), respectively, all assigned hotspot candidate genes are directly related to the biosynthesis of secondary metabolites. Most prominent is the glucosinolate metabolism (Figure 2C), which is represented by three QTL hotspots, explaining variation in 82 metabolites. While variation in various non-volatile intact glucosinolates maps to all three QTL hotspots, variation in the well-known volatile glucosinolate breakdown products, such as nitriles and isothiocyanates (Hansen et al., 2008, Textor et al., 2004, Kliebenstein et al., 2001) maps predominantly to the *AOP* and *MAM* loci. Other QTL hotspots mainly represent phenolic compounds, such as phenylpropanoids and flavonoids (Goujon et al., 2003, Ishihara et al., 2016, Kim et al., 2004, Ross et al., 1999). The analysis of QTL hotspots illustrates that many secondary metabolites are regulated simultaneously by natural variation in a small number of biosynthesis genes upstream of branching points in metabolic pathways. However, enrichment analysis demonstrates that many more genes involved in secondary metabolic processes are assigned as candidate genes explaining variation in non-volatile metabolites than can be expected by chance (43 genes, *P*_*BENJAMINI*_ < 0.01) (Supplemental Table 3, Figure 2B) (Huang et al., 2008). A similar observation was made for candidate genes explaining variation in volatile compounds (Supplemental Table 3, Figure 2B). In addition, many genes assigned to explain variation in specific annotated compounds, after WGR analysis, are directly involved in their biosynthesis pathway (Supplemental Table 4). These results strongly suggest that, although variation in a few genes has a large broad-spectrum effect on metabolic profiles, the widespread natural variation in plant secondary metabolite content is largely driven by variation in a high number of specific biosynthesis genes, consistent with a scale-free model of biological networks. Moreover, these analyses provide statistical evidence that many of the associations detected by WGR mapping are true positives, indicating that variation in a large number of secondary metabolite biosynthesis genes is maintained in nature by selective processes (Bac-Molenaar et al., 2015a).

**Figure 2:**
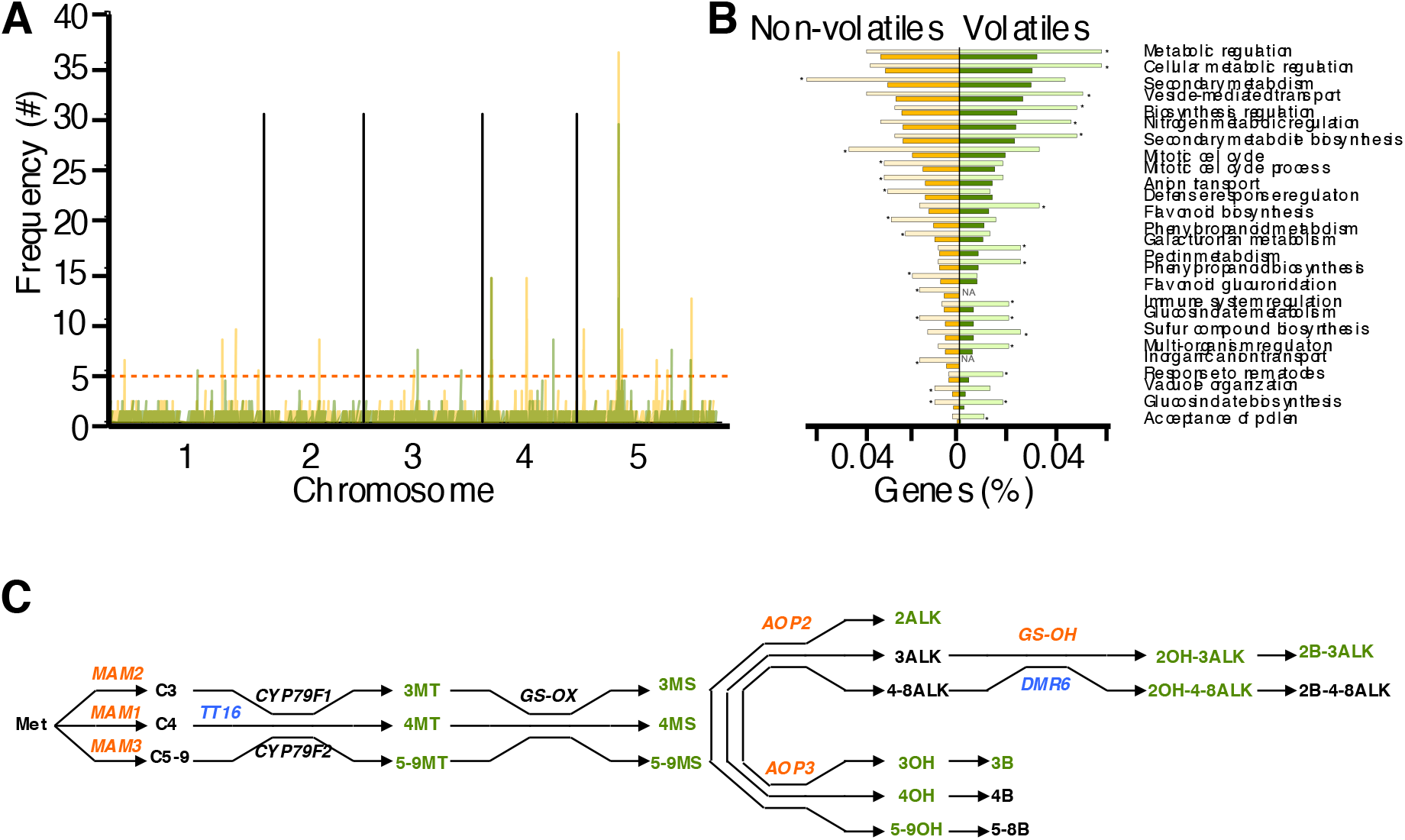
Genetic mapping of metabolic variation. (**A**) Frequency distribution of QTLs over the five chromosomes of Arabidopsis. A QTL hotspot is defined as a locus associated with variation in at least five metabolites. Yellow and green bars indicate the number of non-volatile and volatile metabolites for which a QTL was detected, respectively. The red dotted horizontal line indicates the QTL hotspot threshold. (**B**) Enrichment analysis of candidate genes associated with variation in metabolic content. Dark yellow and green bars indicate the reference genome-wide percentage of genes involved in the processes plotted on the right. Light yellow and green bars indicate this percentage for candidate genes, associated with variation in non-volatile and volatile metabolites, respectively. Asterisks indicate significant enrichment (*P*_*BENJAMINI*_ < 0.01), while *NA* denotes less than two candidate genes involved in the respective process. (**C**) Biosynthesis pathway of glucosinolates. Genes involved in biosynthesis are plotted above arrows. Red genes were assigned as candidate genes explaining the observed variation in glucosinolates. Blue genes have not previously been annotated as biosynthesis genes but were strongly associated with the observed variation in glucosinolates. Synthesised glucosinolates are plotted following arrows. Green glucosinolates were detected and putatively annotated in the current study. C3, C4 and C5-9 indicate glucosinolate carbon-chain-length; MT is Methylthioalkyl; MS is Methylsulfinylalkyl; ALK is Alkenyl; OH is Hydroxyalkyl; OH-ALK is Hydroxyalkenyl; B is Benzoyloxyalkyl; B-ALK is Benzoyloxyalkenyl.

### The role of genetic heterogeneity and inconsistent impact in controlling metabolic diversity

A large advantage of GWR over conventional GWA approaches is the ability to assess more precisely the effect size (*β*) of each SNP as a proportion of the total genetic variance (*σ*^2^_*g*_) explaining quantitative variation in traits. To gain insight into the genetic architecture regulating the variation in metabolic content, SNPs were partitioned into categories of uninformative (*β* = 0*σ*^2^_*g*_), small (|*β*| > 0.0001*σ*^2^_*g*_), intermediate (|*β*| > 0.001*σ*^2^_*g*_) and large (|*β*| > 0.01*σ*^2^_*g*_) effect sizes for each metabolite using GWR (Moser et al., 2015). Although the correct estimation of SNP effect sizes is compromised in GWA mapping by dependency on MAF, a significant correlation was observed between the effect sizes of the most informative SNPs obtained by WGR and GWA for both volatiles (R^2^ = 0.33) and non-volatiles (R^2^ = 0.65) (Supplemental figure 1). Such a relationship was absent when all informative SNPs were included. This is partly due to the strong LD between many SNPs at particular loci and emphasizes the importance of bias reduction, as applied in WGR. Importantly, regardless of the mapping approach followed, the same candidate genes, assumingly causal for the observed variation in metabolite abundance, were generally assigned.

The observed discrepancy in heritability and number of detected QTLs between volatiles and non-volatiles is also reflected in the division of SNP effect sizes. On average, the number and proportion of detected large-effect SNPs per metabolite is higher for non-volatiles (*nV*) than for volatiles (*V*): (*n*_*nV*_ = 1292.8 (|*β*| > 0.0001), 52.8 (|*β*| > 0.001), 3.5 (|*β*| > 0.01) vs. *n*_*V*_ = 1225.4 (|*β*| > 0.0001), 32.9 (|*β*| > 0.001), 1.8 (|*β*| > 0.01) (Supplemental table 2). In total, 52% of the genetically explained variation in non-volatile metabolites is determined by large-effect SNPs, whereas this is only 38% for volatiles. In comparison to other metabolites, the genetically explained variation of metabolites for which a QTL (|*β*| > 0.01) was detected was on average more strongly determined by SNPs of large effect (*nV*: 65% *vs*. 38%; *V*: 56% *vs*. 32%) with higher maximum absolute SNP effect sizes (*nV*: |*β*| = 0.061 *vs*. 0.004; *V*: |*β*| = 0.050 *vs*. 0.006).

However, analogous to the differences in regulation between volatiles and non-volatiles, large inconsistencies do also occur between and within metabolic classes. For instance, 68% of the genetically explained variation in aliphatic glucosinolates is accounted for by large-effect SNPs, whereas this is only 57% and 53% for phenolics and volatile glucosinolate derivatives, respectively. In addition, within the metabolic class of aliphatic glucosinolates, 85% of the genetic variation in short-chain C3-OH glucosinolates is explained by large-effect SNPs, in contrast to 31% for the long-chain C5-C8 MT/MS glucosinolates (Supplemental Figure 2).

The analysis of the regulatory landscape of plant secondary metabolism revealed that many metabolites are regulated by a majority of unique biosynthesis genes and a few hub genes, controlling variation in the abundance of numerous different metabolites. We investigated this observation further by accounting for the effect sizes of SNPs and their associated candidate genes in regulating plant metabolic content. We defined major genes as genes associated with QTLs represented by SNPs with extraordinary high effect sizes (|*β*| > 0.05) and a high likelihood of contributing to the explained variance (Bayes posterior inclusion probability (PIP) > 0.5). As such, 55 major QTLs controlling variation in non-volatiles were detected including eleven of the twelve QTL hotspots (Supplemental Table 5). Similarly, 39 major QTL controlling variation in volatile metabolites were detected, among which the QTL hotspots *AOP* and *MAM* (Supplemental Table 5). Strikingly, non-volatile metabolites (117, *i*.*e*., 20.6%) are much more often regulated by major-effect QTLs than volatiles (35, *i*.*e*., 5.8%). The variation in non-volatile aliphatic glucosinolates and volatile glucosinolate derivatives, for instance, is often determined by one to three QTLs with a major effect and a number of other loci with moderate effects. In contrast, phenolics are mostly regulated by one major QTL and a variety of smaller-effect loci (Supplemental Tables 2, 4 and 6). These analyses highlight the differences in genetic architecture controlling the variation in metabolic content and argues for a power-law distribution of effect sizes in metabolic networks, suggesting that the genetic regulation of metabolic variation leans more towards an additive than an infinite model.

An important topic in the evolution of species is whether adaptive traits have spread through natural populations by drift, recombination and selection of a single alteration or by convergent evolution, in which similar effects are repeatedly obtained by multiple independent mutation events. We addressed this issue by investigating genetic and allelic heterogeneity in the regulation of metabolic content. For this, we performed a detailed analysis of the major-effect WGR QTLs, which are represented by multiple large-effect SNPs (|*β*| > 0.01). Notably, collocating SNPs often differ markedly in effect-size and MAF, and sometimes are not even in LD with each other. This strongly suggests that different haplotypes of particular loci exist within the investigated population, with unequal selection forces acting on the distinctive SNPs. Exemplary, at a number of hotspot QTLs the genetic variation explaining the abundance of a specific metabolite is often determined predominantly by a single major-effect SNP and several smaller-effect SNPs. However, this major-effect SNP is not necessarily the same for other metabolites controlled by the identical hotspot QTL (Supplemental Table 2). So, where independent mutation events at a single locus might lead to allelic heterogeneity in the regulation of one metabolite it might cause haplotype diversity in the regulation of another and, depending on the adaptive value of one or the other metabolite, results in different selective forces on sequence variation.

A different pattern, suggesting alternative modes of regulation, emerges from the analysis of the QTL hotspot at the bottom of chromosome four, explaining variation in 22 non-volatile metabolites. This QTL is represented by two large-effect SNPs in the candidate gene *ACD6*, which are not in LD with each other (R^2^=0.11) (Supplemental Figure 3A). However, the first SNP is in strong LD with various polymorphisms in the *ACD6* promoter, while the second SNP is in strong LD with non-synonymous SNPs in the last exon of the coding region, resulting in the existence of four different haplotypes. Strikingly, 68% of the accessions belonged to haplotype IV, whereas 25%, 6%, and 1% of the investigated accessions belonged to haplotypes III, I, and II, respectively. Haplotype analysis of variation in a representative metabolite controlled by this locus revealed that this metabolite was detected in 90% (18/20) of the accessions of haplotype I, whereas this was only the case for 40% (35/88) and 14% (33/237) of the accessions of the haplotype classes III and IV, respectively. This metabolite could not be detected in any of the accessions (0/2) belonging to haplotype class II (Supplemental Figure 3B). Finally, the abundance of this metabolite was substantially higher in haplotype class I than in any of the other haplotype classes (Supplemental Figure 3C). This indicates an intragenic epistatic interaction effect between the two gene domains. The non-synonymous gene body SNPs might determine the functionality of the gene product and are epistatic over the promoter SNPs, which, in turn, control the level of transcription and, hence, the quantity of the gene product. The variation between haplotypes in the efficiency to produce and accumulate specific metabolites strongly suggests that selective forces have established the observed differences in the global population haplotype frequencies.

Likewise, variation in nine non-volatile metabolites is associated with three SNPs collocating at the *UGT89A2* locus, which together explain up to 37% of the genetic variance in a mixed linear model although none of the SNPs explains more than 24% for any metabolite on their own (Supplemental Table 7). In this case two of the three SNPs exert similar effects on the phenotype, again suggesting allelic heterogeneity, while a specific combination of genotypes is only observed in one accession, suggesting strong negative selection on that haplotype. These analyses indicate that multiple evolutionary events can shape the allelic diversity of genes and might act in concert to obtain distinctive phenotypic levels of metabolites. In addition, it demonstrates that, in these cases, selection acts on haplotypes, determined by multiple SNPs, rather than on single unique SNPs.

In other instances, multiple large-effect SNPs at a single locus are associated to more than one likely candidate gene, suggesting genetic heterogeneity or even channelling of biosynthesis genes (Witjes et al., 2019). The QTL hotspot at the top of chromosome five, explaining variation in the highest number of metabolites, predominantly glucosinolates, links to three genes (*MAM, TT16* and *DMR6*), which all might be involved in metabolite biosynthesis (Falcone Ferreyra et al., 2015, Kroymann et al., 2001, Xu et al., 2017). Variation in the majority of metabolites that map to this locus is best explained by large-effect SNPs that are most strongly associated with *MAM*, although quantitative variation in a minor number of metabolites is stronger linked to large-effect SNPs at the position of *TT16* or *DMR6* (Supplemental Figure 4). Moreover, both *TT16* and *DMR6* are not in strong LD with *MAM* (R^2^ < 0.23). A similar observation can be made for metabolite variation mapping to the QTL hotspot at the top of chromosome four (Supplemental Figure 5). Here, variation in metabolite abundance is most strongly associated to large-effect SNPs at the position of either the *AOP* gene or *DAAR1*, both of which have been implicated in metabolite biosynthesis (Kliebenstein et al., 2001, Strauch et al., 2015). Finally, variation in at least 15 unannotated non-volatile metabolites is associated with no less than seven independent SNPs co-locating at the SAMT/BAMT locus at the lower arm of chromosome five, together explaining up to 57% of the genetic variation in a mixed linear model (Supplemental Table 8). Two of these SNPs are in close proximity and share similar effects on a number of metabolites but display very different MAFs and are not in LD with each other (R^2^ = 0.07), again suggesting allelic heterogeneity. The other five SNPs are more separated and span a region of multiple genes with different metabolic conversion activities, including S-adenosylmethione-dependent methyltransferases (*AT5G37970, AT5G37990* and *AT5G38020*), an UDP-glycosyl transferase (*AT5G38010*), oxidoreductases (*AT5G37940, AT5G37960, AT5G37980* and *AT5G38000*), an arabinosidase (*AT5G37920*) and a general transferase (*AT5G37950*). The metabolites mapping to this locus are not consistent in the SNPs with the strongest associations and thus might be regulated by different genes in this region.

These results illustrate that WGR mapping can be instrumental in disentangling the genetic architecture of metabolic trait regulation by providing a higher resolution of assigning candidate genes to detected genomic associations, through inclusion of accurate SNP effect-size estimates.

### Polygenic adaptation of plant secondary metabolism to local settings

Evolutionary theory predicts that selective forces shift allele frequencies of non-neutral genetic variants in admixed populations (Walsh and Lynch, 2018) and there is no reason to expect other effects on natural variation in secondary metabolism. Indeed, a positive and significant correlation was observed between the BayesR effect-size |β| and MAF of the SNPs most strongly associated (|β|> 0.01 and PIP_large_ > 0.5) with variation in the accumulation of both volatile (R^2^=0.32) and non-volatile (R^2^=0.20) secondary metabolites (Supplemental Figure 6). Such a correlation was absent when all, mostly neutral, SNPs were taken into account, suggesting that balancing selection is a strong driver of maintaining natural variation in secondary metabolism (Benderoth et al., 2006). Following this reasoning it is plausible to assume local adaptation, which should be reflected in the fixation index (F_ST_) of selective loci (Holsinger and Weir, 2009). As expected, SNP F_ST_ values calculated for five European regions correlate positively with the effect-size of the SNPs most informative (|β|> 0.01 and PIP_large_ > 0.5) for variation in volatile and non-volatile metabolites (R^2^=0.42 and 0.22, respectively) (Supplemental Figure 7). Illustratively, approximately 4% of all investigated SNPs pass the F_ST_ threshold value of 0.2, generally considered as indicative for selection, while almost 13% of the SNPs with an effect-size |β|> 0.01 and PIP > 0.5, explaining variation in non-volatile abundance, meet this criterion. This proportion was even higher for SNPs explaining variation in volatiles (21.9%), although the number of qualifying SNPs was much lower. Moreover, a strong enrichment (*P* < 0.005) of 160 genes involved in secondary metabolism was observed among the list of candidate genes assigned to SNPs with F_ST_ values above 0.2, compared to the 132 genes expected by chance. In addition, strong selection (0.21< F_ST_ <0.44) was observed for half the number of QTL-hotspots, including all three loci explaining variation in glucosinolate metabolism, with large-effect SNPs in the 0.05‰ extreme end of the F_ST_ distribution (F_ST_ > 0.3). Other loci explaining variation in glucosinolate, phenylpropanoid, and flavonoid accumulation also displayed high F_ST_ values (Supplemental table 9). Together, these analyses illustrate that local adaptation is an important driver for the evolution of natural variation in secondary metabolism.

The high F_ST_ values observed for some SNPs suggests the existence of population structure and a relationship between metabolic profiles and the geographic origin of accessions. To investigate this further a two-dimensional hierarchical clustering was performed on the geographic distribution of analyzed accessions and the accumulation of detected compounds therein. Not surprisingly, given their strong enrichment in the phytochemical profile, aliphatic glucosinolates and their volatile derivatives were among the strongest determinants of genotype clustering (Supplemental Figure 8A). The aliphatic glucosinolates were further categorized on chain length and side-chain modification and based on this classification four main groups of genotypes, each with a different organization of glucosinolate biosynthesis alleles and enriched in accessions from specific geographic regions, could be distinguished (Supplemental Figure 8B). For instance, alkenyl and long-chain glucosinolates group closely together and are, on average, detected at high levels in the majority of genotypes from the UK and France (group 1). From the 105 accessions within this group, 45% is of French origin (*i*.*e*., 77% of all French accessions) while 27% originates from the UK (*i*.*e*., 60% of all accessions from the UK) (Supplemental Table 1). The qualitative and quantitative variation in glucosinolate content is largely explained by allelic variation in five biosynthesis genes (*viz. MAM, AOP, TT16, DMR6*, and *GS-OH*). Both the *TT16* locus and the *DMR6* locus have not been reported earlier in glucosinolate studies, probably due to the close vicinity to *MAM*. Both loci are, however, not in very strong LD with *MAM* (R^2^ < 0.23). A candidate gene for *TT16* might be coding for a succinyl CoA ligase involved in the TCA cycle (AT5G23250), or a nicotinamidase involved in coniferin metabolism (AT5G23220/30). The *DMR6* gene is co-expressed with *GS-OH*, shares protein domains with *AOP* and *GS-OH* and exhibits, like *AOP* and *GS*-*OH*, oxidoreductase activity (Warde-Farley et al., 2010). Given the variation in C4-8 alkenyl glucosinolates explained by *DMR6*, the loss of an active *GS-OH* allele might be compensated by a similar function of *DMR6*. To test this hypothesis, a population of doubled haploid (DH) lines was generated from a cross between the accessions UKNW06-460 and Tamm-2 and subjected to metabolic analyses. Unlike the reference accession Col-0, which lacks expression of functional AOP (Kliebenstein et al., 2001), these accessions share functional *AOP* and *MAM* genes, necessary for the production of 3-butenyl, which is the precursor for the synthesis of 2-hydroxy-3-butenyl. However, UKNW06-460 and Tamm-2 differ in functionality of the *GSOH* and *DMR6* loci. Tamm-2 lacks functional alleles of both *GSOH* and *DMR6* and is unable to synthesise 2-hydroxy-3-butenyl despite the fair amount of available 3-butenyl, whereas UKNW06-460 contains both a functional *GSOH* and *DMR6* allele, which allows it to produce 2-hydroxy-3-butenyl in addition to 3-butenyl (Supplemental figure 9A). Recombinant DH lines were divided in four haplotype classes based on functionality of the *GSOH* and *DMR6* alleles. With a few exceptions, haplotypes lacking a functional *DMR6* allele (haplotype I and II) did not produce detectable levels of 3-butenyl and 2-hydroxy-3-butenyl. Haplotypes containing a functional allele of *DMR6* (haplotype III and IV), however, were able to synthesise high levels of both compounds irrespective of the presence of a functional *GSOH* allele (haplotype IV) or not (Haplotype III), although the production of 2-hydroxy-3-butenyl is much higher when both genes are functional (Supplemental figure 9B). While the model of redundant functions of the *GSOH* and *DMR6* gene does not unambiguously explain the observed metabolic profiles in natural accessions and DH lines, possibly due to additional segregating genetic modifiers, metabolic feedback, genotype mis-calling or metabolite mis-annotation, the results clearly indicate similarities in function of the *GSOH* and *DMR6* genes.

Group 1 is largely represented by haplotypes containing the Columbia (Col-0) reference allele for *MAM* and *AOP* and the non-Col-0 *TT16* allele, combined with either the non-Col-0 *GS-OH* or *DMR6* allele (Figure 3). A second group consists mainly of accessions from Sweden and the USA, of which the majority contain high levels of C3 and C4 alkenyl glucosinolates. The accumulation of these glucosinolates is determined by a cooperation of the *AOP* and *MAM* loci and haplotypes carrying the Col-0 *AOP* allele and the non-Col-0 *MAM* allele, synthesizing the highest levels of alkenyl glucosinolates, dominate this group (Figure 3). Two other, North-East European, groups are characterized by an overrepresentation of accessions from the Czech Republic (group 3) and Germany (group 4), respectively. Haplotypes in the Czech group share the non-Col-0 *AOP* allele, which is associated with high levels of C3-hydroxyl glucosinolates, while haplotypes in the German group share the Col *TT16* allele and produce large amounts of C4-methylthio/methylsulfinyl glucosinolates (Figure 3). These analyses illustrate that glucosinolate profiles are strongly determined by local adaptive processes that have shaped the allelic diversity at various biosynthesis loci. These loci explain on average a large proportion (41%) of the observed variation in aliphatic glucosinolates (Supplemental Table 10) and, depending on the specific haplotype, result in a semi-qualitative distribution in almost discrete classes of accumulation (Figure 3). For most aliphatic glucosinolates this variation is almost completely explained by additive effects of contributing loci, while epistatic interactions play a very minor role (Supplemental Table 10).

**Figure 3:**
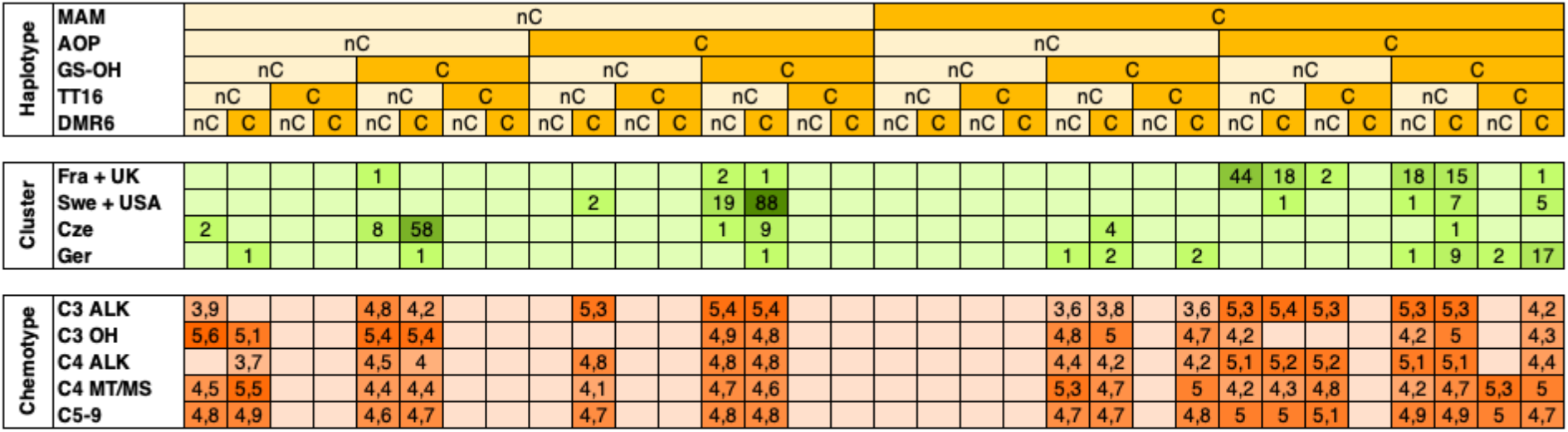
Genotypic and geographic patterns of adaptation of glucosinolate profiles. The upper panel divides haplotypes based on the genotypes of five loci involved in the biosynthesis of glucosinolates. C denotes Col-0 alleles, while nC denotes non-Col-0 alleles. The second panel divides the collection of Arabidopsis accessions analyzed in three geographical clusters based on hierarchical clustering of metabolite profiles. Columns align with the observed haplotype indicated in the upper panel. For each cluster and haplotype the number of accessions is given. The lower panel divides the detected glucosinolates in five classes. Columns align again with the observed haplotype indicated in the upper panel. For each haplotype the relative average abundance of glucosinolates assigned to a specific chemotype is given.

Additive effects increase the selection coefficient of specific loci, which drives the co-evolution of certain allele combinations in particular environments, as was previously observed for glucosinolate metabolism (Brachi et al., 2015). In line with this, the occurrence of some haplotypes deviates significantly from estimates based on population allele frequencies. For instance, the non-Col-0 *GS-OH* allele almost exclusively occurs together with the Col-0 *MAM* and *AOP* allele, while the combination of the nonCol-0 *AOP* allele and the Col-0 *MAM* allele is extremely rare (Figure 3). Interestingly, a sixth locus, encoding an epithiospecifier protein (*ESP*) involved in glucosinolate breakdown and explaining a large fraction of the variation in C4-8 alkenyl glucosinolates acts hypostatic to the *GS-OH* and *MAM* loci (Supplemental Table 11). A co-evolution of *ESP* with *GS-OH* and *MAM*, and to a lesser extent *DMR6*, is further suggested by moderate to high levels of LD between SNPs representing the various loci, even though located on different chromosomes (Supplemental Figure 10). The active non*Col-0* ESP allele is found in all, but one accessions from France and the UK (Supplemental Table 12), suggesting a favourable production of nitriles over isothiocyanates in these regions. In contrast, the vast majority of accessions from the Czech Republic carry the weak *Col-0* allele, further confirming strong population structure for the accumulation of these compounds.

Analogous to the glucosinolates and their volatile derivatives many other metabolites, including phenylpropanoids and flavonoids, display strong correlations with geographical clines and climate parameters (Supplemental Table 13). In addition, the loci explaining most of this variation, like *BGLU6* and *UGT78D2*, exhibit high F_ST_ values and a strong correlation of allelic variation with latitude and longitude (Supplemental Tables 9 and 13).

This study demonstrates that natural variation in plant secondary metabolism is governed by allelic variation in many biosynthesis genes, of which the effect size could be accurately determined by Bayesian statistics. A substantial part of the allelic diversity is likely shaped by local adaptation to resident climates with strong selection for specific haplotypes and metabolic profiles.

## Discussion

In this study we provide evidence of selective forces acting on adaptive allelic variation in biosynthesis genes of secondary metabolites, shaping regional variation in phytochemical profiles. Moreover, we were able to estimate accurately the effect size of genetic variants and establish the genetic architecture of the regulatory landscape of plant secondary metabolism. Using state-of-the-art untargeted metabolomics more than a thousand volatile and non-volatile secondary metabolites could be comparatively quantified in a global collection of more than 350 Arabidopsis accessions. For most of these compounds, heritable qualitative and quantitative variation was observed and for approximately half of the non-volatile, and a quarter of the volatile metabolites, this variation could be explained by genetic variation in specific genomic loci, according to a Bayesian WGR approach. A majority of these loci could be tied to candidate genes involved in the biosynthesis pathway of the metabolite under study, suggesting that most of the identified associations are true positives. Furthermore, many metabolites mapped to the same genomic loci, resulting in a total of ten hotspots explaining variation in no less than 44 and 120 volatile and non-volatile metabolites, respectively. Surprisingly, very little overlap was observed in QTL hotspots and candidate genes assumed to be causal for the observed variation in volatile and non-volatile metabolites. This suggests an independent regulation of these two types of compounds, which can be explained by regulation *in cis* of key biosynthesis genes of specific metabolites or modules rather than a systematic overarching regulation *in trans* of whole plant secondary metabolism.

The WGR enabled estimation of the effect-size of genetic variation allowed the classification of contributing loci and a more detailed analysis of specific loci. Apparently, one-third to half of the secondary metabolites is regulated by large-effect polymorphisms, although large differences occur between and within metabolic classes. This supports the generally accepted view that the genetic basis of plant secondary metabolism is founded on a few large-effect loci, numerous small effect-loci, and genetic and environmental interactions (Olson-Manning et al., 2012). Interestingly, on several loci, multiple independent mutation events could be detected, resulting in a variety of different haplotypes in the global Arabidopsis population. Although allelic or genetic heterogeneity might occur, signatures of selection indicate that, depending on the geographic origin of the accessions, unequal selective forces act on the uncovered genetic variation, resulting in different effects on the accumulation of metabolites. These findings correspond well with the observed dependency of fitness effects of metabolic profiles on field conditions (Kerwin et al., 2015). Indeed, a positive correlation was detected between the effect-size, F_ST_ value and allele frequency of genetic variation and a strong enrichment of QTL hotspots and candidate genes involved in secondary metabolism with high F_ST_ values. The data presented here support the hypothesis that natural variation in secondary metabolism is maintained by fluctuating or balancing selection throughout the long evolutionary history of Arabidopsis (Kliebenstein et al., 2016).

The relationship between the geographic origin of accessions and their metabolic profile can, at least for aliphatic glucosinolates, largely be explained by additive effects of genetic variation. This is consistent with reports of allelic variation for morphological and physiological traits being correlated with climate and geographical variables (Baxter et al., 2010, Dubin et al., 2015, Hancock et al., 2011, Kooke et al., 2016, Li et al., 2010). In addition, the observed frequency of haplotypes differs from expectations of neutral theory, indicating local adaptation and selection of favourable combinations of alleles of different genes. Moreover, significant LD could be detected between loci involved in the joint regulation of specific metabolites, even when these are located on different chromosomes, speculating on the co-evolution of genes establishing biosynthesis pathways. Co-evolution and epistatic selection have previously been suggested for important loci controlling glucosinolate metabolism, such as *GS-OH, AOP* and *MAM* (Brachi et al., 2015), and our data suggest that other genes, such as *DMR6, ESP* and *CYP79F2*, have undergone similar selective processes and might have an impact on fitness as well. Interestingly, recent analyses indicate a role for epigenetic regulation of metabolic content as well, which, in contrast to the irreversible adjustments described here, might provide a more flexible adaptation to alternating conditions (Kooke et al., 2019).

Our results provide compelling additional evidence that natural variation in the accumulation of secondary metabolites in *Arabidopsis thaliana* is governed by local adaptation through evolutionary selection of polygenic variation.

## Material and Methods

### Plant growth conditions

Seeds of 359 Arabidopsis natural accessions belonging to the HapMap panel (Horton et al., 2012, Li et al., 2010) were sown on filter paper with demi water, stratified at 4°C in dark conditions for five days and transferred to a culture room (16h LD, 24°C) for 42h to induce seed germination. Eight replicates per accession were transplanted to wet Rockwool blocks of 4×4 cm in a completely randomized block design in a climate chamber. Chamber climate conditions were as follows: 12 h short days, light intensity 125 µmol.m^-2^.s^-1^, temperature 20°C day/18°C night, relative humidity 70%. All plants were watered daily for five minutes with 1/1000 Hyponex solution (Hyponex, Osaka, Japan). Two replicates of each accession were harvested in bulk to serve as reference material in metabolite analyses. The other plants were harvested in two replicate pools of three plants 28 days after sowing. The harvest time was set to the end of the light period.

### Metabolomics

Samples were ground in liquid nitrogen and an aliquot of all samples was mixed to generate a large pool needed for preparing the quality controls (QCs). These were independently and simultaneously weighed and extracted with the study samples (5–6 times per batch) and injected at regular intervals within the analysis series.

For the detection of non-volatiles, aqueous-methanol extracts were prepared from 50 mg frozen ground material to which 200 μl of 94 % MeOH containing 0.125 % formic acid was added (De Vos et al., 2007). Batch sizes ranged from 78 to 80 samples, with the exception of the last batch, batch 10, containing 48 samples (Wehrens et al., 2016). After sonication and filtering, the crude extracts were analysed as described previously (van Duynhoven et al., 2014) using UPLC (Waters Aquity) coupled to a high-resolution Orbitrap FTMS (Thermo). A 20 min gradient of 5–35 % acetonitril, acidified with 0.1 % formic acid, at a flow rate of 400 μl/min was used to separate compounds on a 2.1 × 150 mm^2^ C18-BEH column (1.7 μm particle size) at 40 °C. Metabolites were detected using a LTQ-Orbitrap hybrid MS system operating in negative electrospray ionization mode heated at 300 °C with a source voltage of 4.5 kV. The transfer tube in the ion source was replaced and the FTMS recalibrated after each sample batch, without stopping the UPLC system. After pre-processing raw data files in an untargeted manner using a Metalign-MSClust based workflow (Kooke et al., 2019, Lommen and Kools, 2012, Tikunov et al., 2012), metabolites occurring in fewer than 20 different genotypes were removed, leading to a data matrix containing relative intensities of 567 reconstructed metabolites in 761 samples (including 51 QCs). The percentage of non-detects (*i*.*e*., an intensity value below the arbitrarily set detection threshold) in this matrix is 48 %. For individual metabolites, the fraction of non-detects can be much larger, and in this data set is up to 97 %.

The detection of volatiles is based on aliquots of the same Arabidopsis material as described for the non-volatiles. The aim here was to analyse volatile organic compounds (VOCs) present in the leaf material using solid phase microextraction (SPME) of the headspace. Extracts of 50 mg from frozen ground material were analysed on a GC–MS system (Agilent GC7890A with a quadrupole MSD Agilent 5978C) in fifteen batches of 34-99 samples, with, on average, 15 study samples per QC, as described earlier (Cordovez et al., 2015, Mumm et al., 2016, Verhoeven et al., 2012). In contrast to these studies, the temperature program of the GC oven started at 45 °C (2 min hold) and rose first with 8–190 °C.min^-1^, followed by 25–280 °C (2 min hold). This data set contains information on 753 samples (including 50 QCs) with, in total, 40 % non-detects, similar to what was found for the non-volatiles. For individual metabolites, the percentage of non-detects goes up to 97 %. As for the non-volatiles, only those volatile metabolites were retained that were present in at least 20 different genotypes, in this case 603 metabolites.

In all subsequent analysis, log-transformed metabolite intensity values were used. Accessions in which the relative abundance of the assayed metabolite did not pass the detection threshold were assigned the threshold value of 3.0 and 0.45 for non-volatiles and volatiles, respectively.

### Descriptive statistics

The variance components for all the individual traits were used to calculate the broad-sense heritability, *H*^*2*^, in analysis of variance (ANOVA) according to the formula *H*^*2*^ *= σ*^*2*^_*G*_*/ (σ*^*2*^_*G*_ *+ σ*^*2*^_*E*_), with *σ*^*2*^_*G*_ *= (MS(G) – MS(E))/r, σ*^*2*^_*E*_ *= MS(E)*, where *r* is the number of replicates and *MS(G)* and *MS(E)* are the mean sums of squares for genotype and residual error, respectively. Narrow-sense heritability, *h*^*2*^, was defined as *h*^*2*^ *= σ*^*2*^_*A*_*/(σ*^*2*^_*G*_ *+ σ*^*2*^_*E*_), which takes only the additive genetic effects (*σ*^*2*^_*A*_) in account. Marker-based estimates of narrow-sense heritability were obtained from the mixed model, *y*_*i,j*_ *= μ + G*_*i*_ *+E*_*i,j*_, *(i = 1,…,n, j = 1,…*.,*r), G* ∼ *N(0, σ*^*2*^_*A*_*K), E_i,j ∼ N(0, σ*^*2*^_*E*_), where *y*_*i,j*_ is the phenotypic response of replicate *j* of genotype *i, μ* is the intercept, *G = (G*_*1*_,*…,G*_*n*_) is the vector of random genetic effects, and the errors *E*_*i,j*_ have independent normal distributions with variance *σ*^*2*^_*E*_ which is the residual variance for a single individual (Kruijer et al., 2014). The vector *G* has a multivariate *N(0, σ*^*2*^_*A*_*K)* distribution, and the genetic relatedness matrix *K* is estimated from standardized SNP-scores. The coefficient of variation (*CV*_*G*_) was calculated as 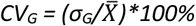.

### Genome-wide association mapping

All accessions were genotyped with 214,051 SNPs (Li et al., 2010) of which, after removal of SNPs with a minor allele frequency (MAF) < 0.05, 199,589 were used for genome wide association mapping.

All traits were analysed with a Bayesian statistical model, BayesR, which uses a Gibbs sampling approach to estimate variant effects that are modelled as a mixture distribution of four normal distributions. SNPs were assigned a prior variance of 0*σ*^2^_*g*_, 0.0001*σ*^2^_*g*_, 0.001*σ*^2^_*g*_, and 0.01*σ*^2^_*g*_, across the four distributions, respectively, where *σ*^2^_*g*_ is the total genetic variance. This allows many uninformative variants to be dropped from the model and permits remaining variants to have moderate to large effects. Gibbs sampling was performed for 50,000 iterations, after 20,000 burn-in iterations. Conventional GWAS was performed using the R-package statgenGWAS (https://github.com/Biometris/statgenGWAS/), following the approach of previous studies (Kang et al., 2010, Kruijer et al., 2014).

### Enrichment analysis

Enrichment analysis was performed using the functional annotation tool in DAVID 6.8 (https://david.ncifcrf.gov/home.jsp) (Huang et al., 2008) for the candidate gene lists of the volatile and non-volatile compounds separately. The gene lists were run against a background set of Arabidopsis TAIR IDs. The gene ontology category GOTERM_BP_FAT was explored and the significance threshold for enrichment was set at *P* < 0.01.

### Fixation index

A large subset of the global Arabidopsis population was divided into five sub-populations based on the countries of origin that were best represented in the collection, *viz*. Germany (57 accessions), France (56 accessions), Sweden (48 accessions), UK (47 accessions), and the Czech Republic (29 accessions). These accessions together comprised 68% (237 out of 350 genotypes) of the entire population. F_ST_ values were calculated using the following formula: *F*_*ST*_ *= (π*_*Between*_ *– π*_*Within*_*)/π*_*between*_, where *π*_*Between*_ and *π*_*Within*_ represent the expected heterozygosity (*i*.*e*., genetic diversity) between populations and the expected heterozygosity within populations, respectively.

### Validation of *GSOH*/*DMR6* redundancy

To test for redundancy of the *GSOH* and *DMR6* gene functions three accessions were selected that differed in functional alleles for these two genes and *AOP*. Col-0 (CS76113) does not express functional alleles of *AOP* and is unable to synthesize 3-butenyl, the precursor for the synthesis of 2-hydroxy-3-butenyl (Kliebenstein et al., 2001). UKNW06-460 (CS76279) carries both a functional *GSOH* and *DMR6* gene, whereas Tamm-2 (CS76244) lacks functional alleles for both genes. From a cross between Tamm-2 (♀) and UKNW06-460 (♂) 88 double haploid lines were generated (Filiault et al., 2017), which were grown in a climate-controlled growth chamber for 28 days in short-day conditions (12 h, 125 µM, 70% RH, 20/18°CD/N). After harvesting, plants were snap-freezed and stored in -80°Cupon further analysis. Subsequently, all DH lines were genotyped with KASPar™ genotyping technology (KBiosciences), distinguishing the parental haplotype for *GSOH* and *DMR6* at two SNP-positions for each gene. Primers were developed on the following sequences: GS-OH_1 (Chr2: 10799538 bp) CS76244/CS76279 > gagagacttctcaattcaaaaacagacatggaggatctta/gtagcgagactaaatcaagagacagcggtgaaggaatatc; GS-OH_2 (Chr2: 10832548 bp) CS76244/CS76279 tgtcaggatcaaattcaatttcaataaccagtcacaatatctttcaatta/tctggaatttctatctacatttagcataacctttcatattattttttacac; > DMR6_1 (Chr5: 8379646) CS76244/CS76279 > atctaagactccattgttatcctatccacaagtatgtcc/aatgagtggccgtcaaaccctccttctttcaagtaagca; DMR6_2 (Chr5: 8366765) CS76244/CS76279 > gttggtccagacttacctaataggctaatgcctta/ttcattggagtttgctgatcttatgatgacgtttt, according to standard protocol (Smith and Maughan, 2015). DH lines were classified in one of four different genotypic classes (haplotype I: *GSOH* inactive/*DMR6* inactive; haplotype II: *GSOH* active/*DMR6* inactive; haplotype III: *GSOH* inactive/*DMR6* active; haplotype IV: *GSOH* active/*DMR6* active. For each haplotype three independent pools of two plants each (*i*.*e*., 3 × 2 different DH lines) were collected from which a single methanol extract was obtained as described above. Extracts were then subjected to UPLC FTMS as described above and accumulation of 3-butenyl and 2-hydroxy-3-butenyl was quantified as the percentage of detection saturation.

### Climate Data

Climate data for the collection origin of each accession was obtained from the Climate Research Unit at the University of East Anglia. Data were extracted for nine climate variables giving the average per month over a 30-year (1961-1990) period (New et al., 2002). From these 9 variables, most other variables were extracted. Day length (spring) and relative humidity (spring) from the site of 306 accessions were obtained from the NCEP-NCAR climate reanalysis project (Hancock et al., 2011, Kistler et al., 2001) and the FAO GeoNetwork (http://fao.org.geonetworks/srv/en/main.home).

## Acknowledgements

RK was funded by grants from CBSG (AA3-WU-PL) and STW (STWGrant 1099).

