## Supplemental figures for "Local adaptation shapes metabolic diversity in the global population of *Arabidopsis thaliana*"

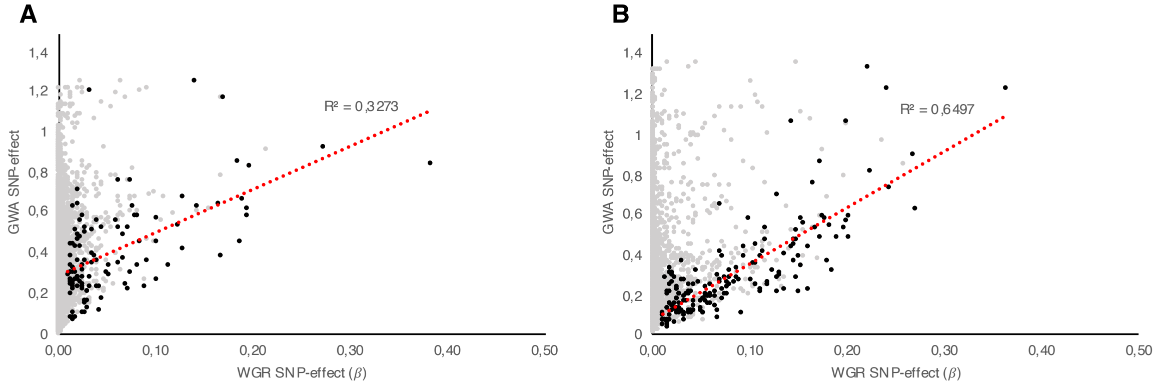


**Supplemental figure 1:** Correlation between SNP effect sizes estimated by Bayesian WGR and EMMAX GWAS. (**A**) Correlation between effect sizes of SNPs associated with variation in volatile metabolites. (**B**) Correlation between effect sizes of SNPs associated with variation in non-volatile metabolites. Black dots represent large-effect SNPs (β > 0.01), for each metabolite only the largest-effect SNP was included. The red dotted line depicts the regression of the data. Grey dots represent all other informative SNPs as determined by Bayesian WGR.

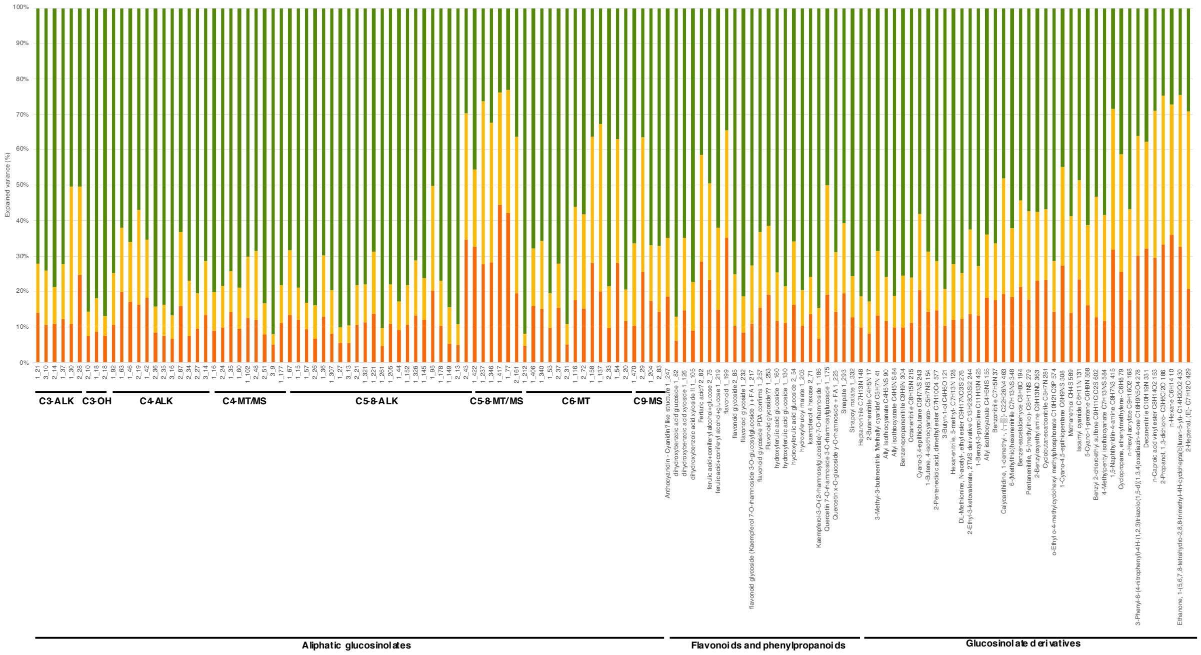

**Supplemental figure 2:** Genetic architecture of the regulation of metabolite accumulation.
Plotted is the percentage of genetic variation explained by large-effect (green), medium-effect (yellow) and small-effect (red) SNPs. Non-volatile aliphatic glucosinolates are subdivided in metabolic classes according to chain length and side-chain modification. Non-volatile flavonoids and phenylpropanoids are indicated by IUPAC name or synonym. Volatile glucosinolate derivatives are indicated by IUPAC name and molecular formula. All compounds are also indicated by the mass-ID from this study.

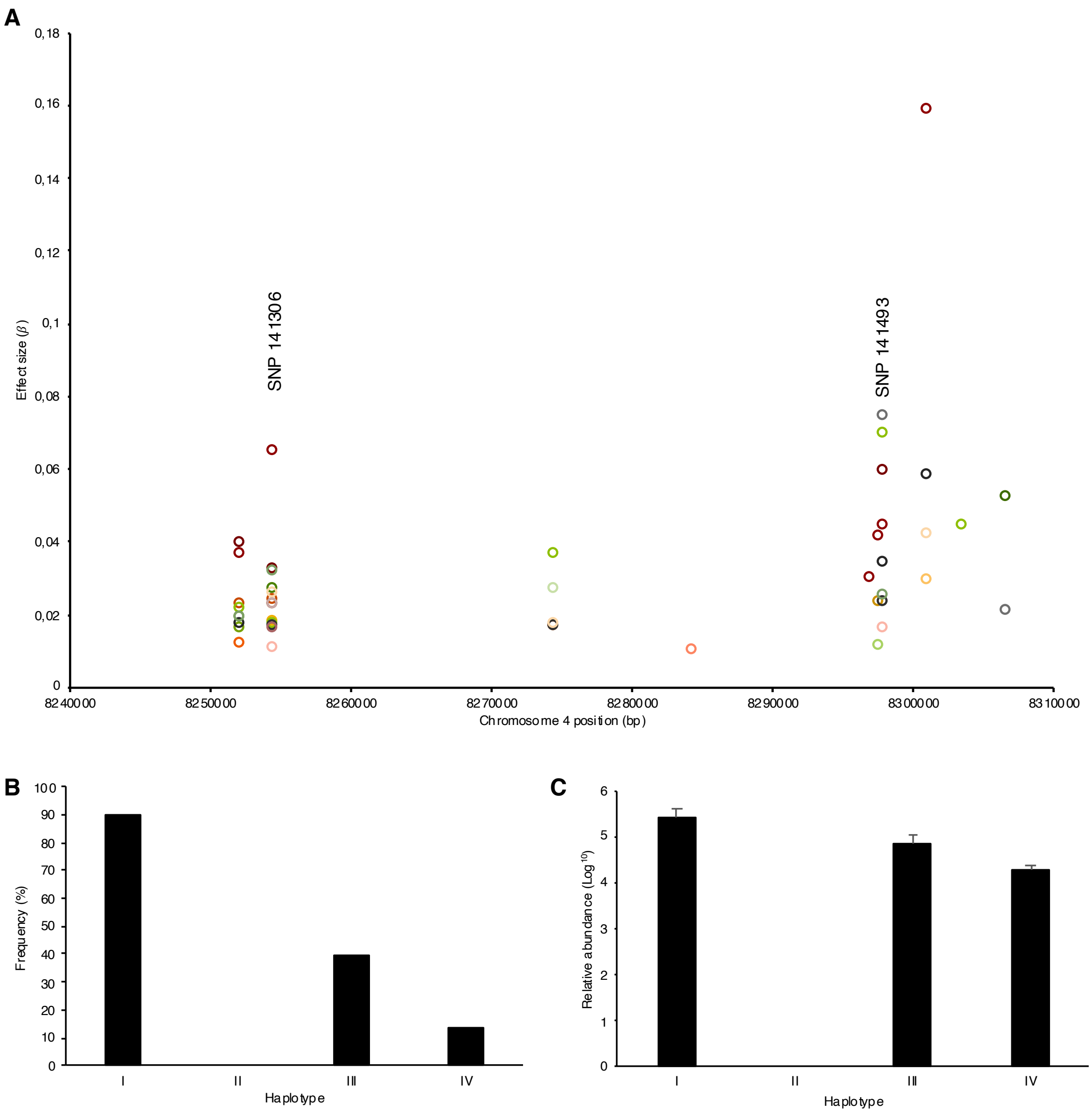


**Supplemental figure 3:** Intragenic epistasis between genetic variants at the *ACD6* locus.
(**A**) Effect sizes of SNPs in the vicinity of the ACD6 gene. Each color of the open circles represents the major effect size of a SNP at that position for one of 22 different metabolites. The position of two large-effect SNPs in the promoter region (SNP 141306; MAF 0.94) and gene body (SNP 141493; MAF 0.31) are indicated. (**B**) Occurrence of the representative but unknown non-volatile metabolite 1-354 in different haplotypes of the Arabidopsis population. The haplotype is determined by the genotype of the two SNPs indicated in figure 3A (Haplotype I: SNP 141306 = Col-0, SNP 141493 = Col-0; Haplotype II: SNP 141306 = Col-0, SNP 141493 = nonCol-0; Haplotype III: SNP 141306 = nonCol-0, SNP 141493 = Col-0; Haplotype IV: SNP 141306 = nonCol-0, SNP 141493 = nonCol-0). (**C**) Relative abundance of metabolite 1-354 in accessions in which this metabolite was detected for the different haplotype classes. Note the log-scale on the y-axis. Error bars represent standard error of the mean (SEM).

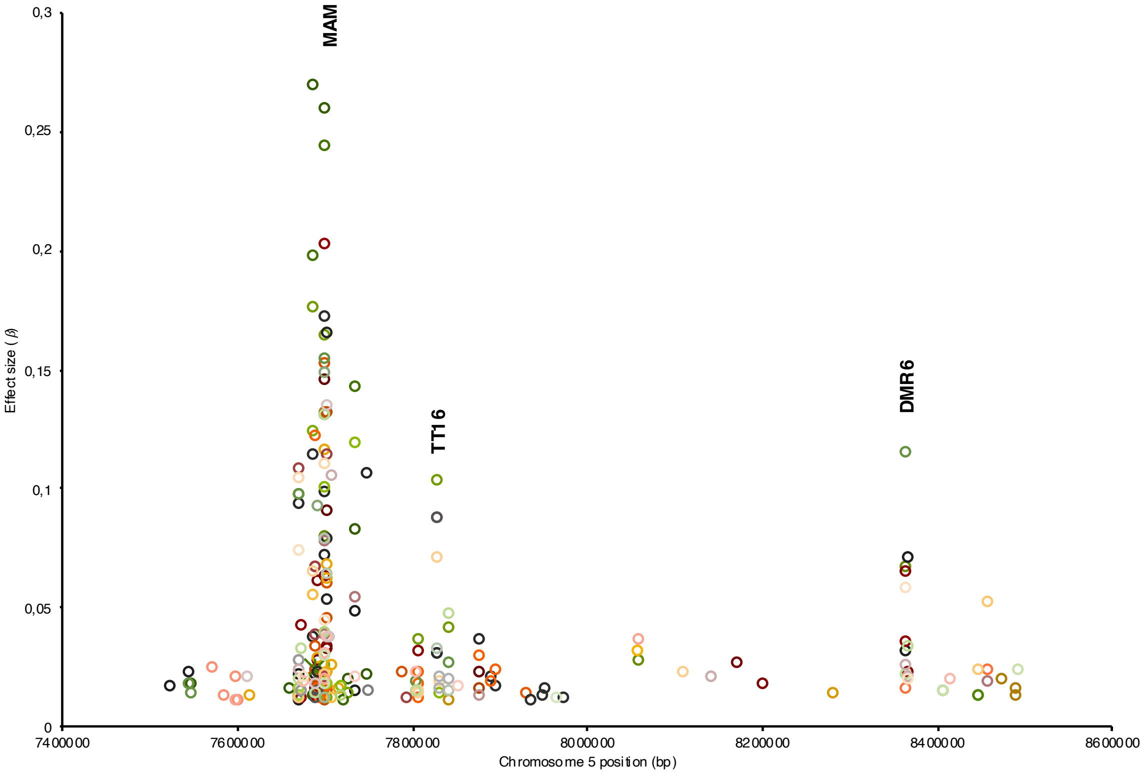


**Supplemental figure 4:** Genetic heterogeneity at the MAM QTL hotspot.
Effect sizes of SNPs in the vicinity of the *MAM* gene are plotted. Each color of the open circles represents the major effect size of a SNP at that position for one of 58 different metabolites. The position of the candidate genes MAM, TT16 and DMR6 are indicated in bold face.

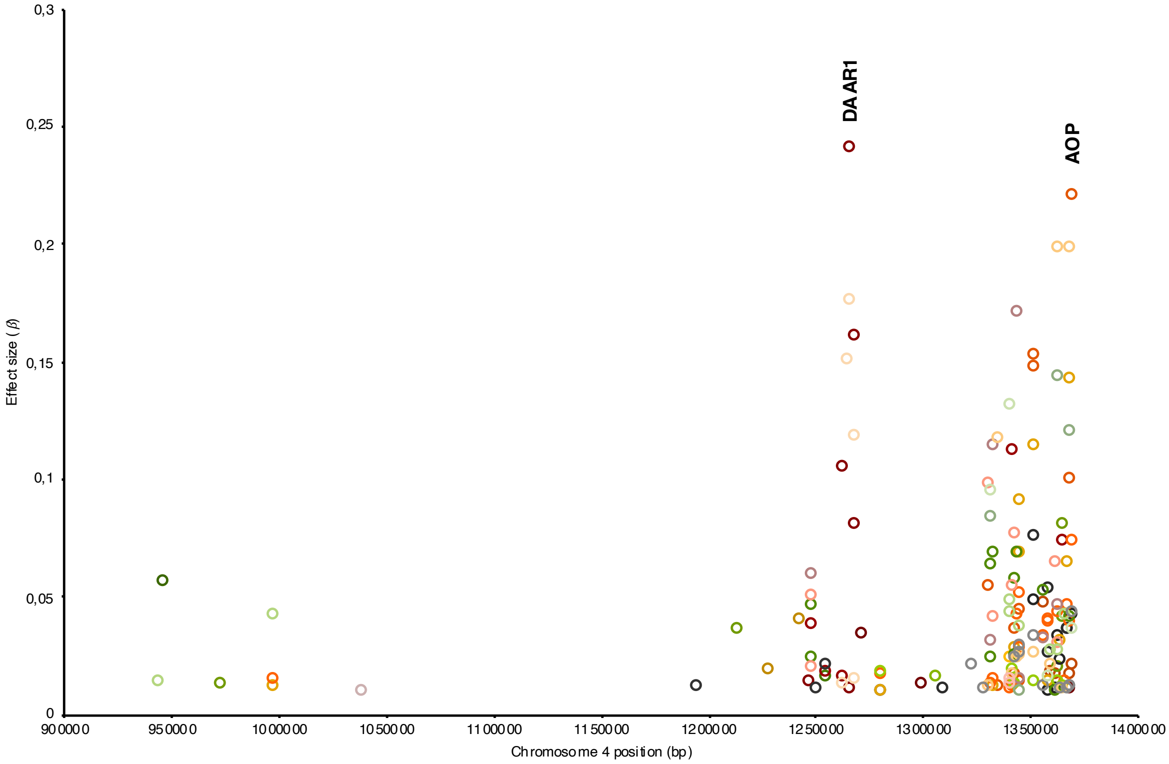


**Supplemental figure 5:** Genetic heterogeneity at the AOP QTL hotspot.

Effect sizes of SNPs in the vicinity of the *AOP* gene are plotted. Each color of the open circles represents the major effect size of a SNP at that position for one of 28 different metabolites. The position of the candidate genes *DAAR1* and *AOP* are indicated in bold face.

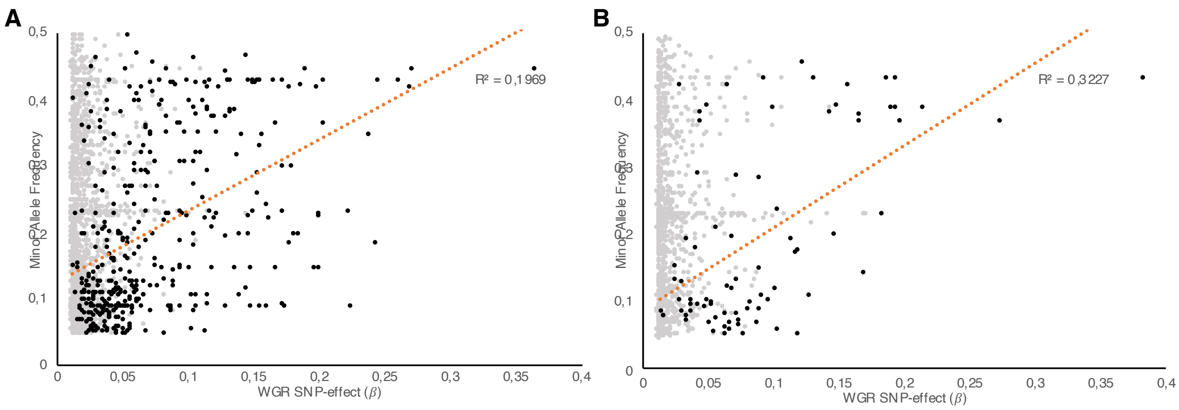


**Supplemental figure 6:** Correlation between SNP effect-size and minor allele frequency.
(**A**) Correlation between effect size and MAF of SNPs associated with variation in non-volatile metabolites. (**B**) Correlation between effect size and MAF of SNPs associated with variation in volatile metabolites. Black dots represent the largest-effect SNPs (β > 0.01, PIP > 0.5), for each metabolite only one SNP was included. The red dotted line depicts the regression of the data. Grey dots represent all other informative SNPs as determined by Bayesian WGR (β > 0.01, PIP < 0.5).

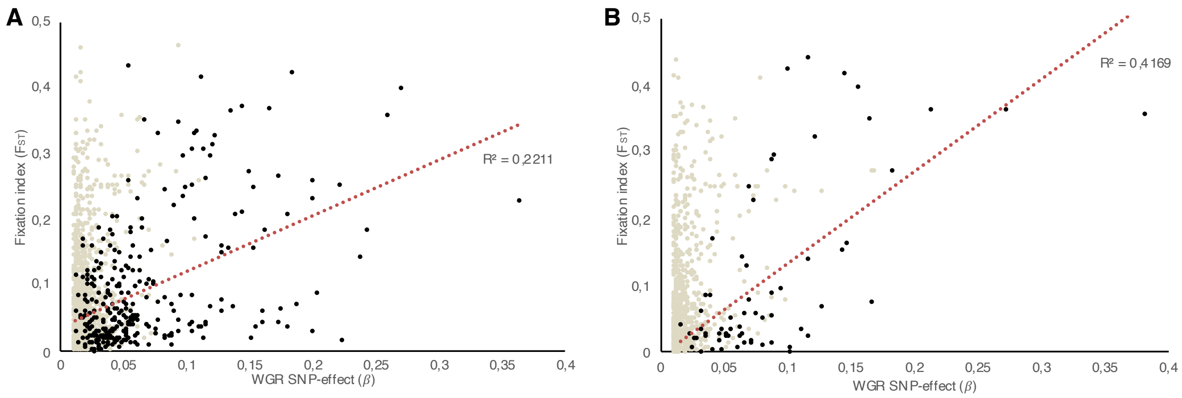


**Supplemental figure 7:** Correlation between SNP effect-size and fixation index.
(**A**) Correlation between effect size and $F_{ST}$ of SNPs associated with variation in non-volatile metabolites. (**B**) Correlation between effect size and $F_{ST}$ of SNPs associated with variation in volatile metabolites. Black dots represent the largest-effect SNPs (β > 0.01, PIP > 0.5), for each metabolite only one SNP was included. The red dotted line depicts the regression of the data. Grey dots represent all other informative SNPs as determined by Bayesian WGR (β > 0.01, PIP < 0.5).

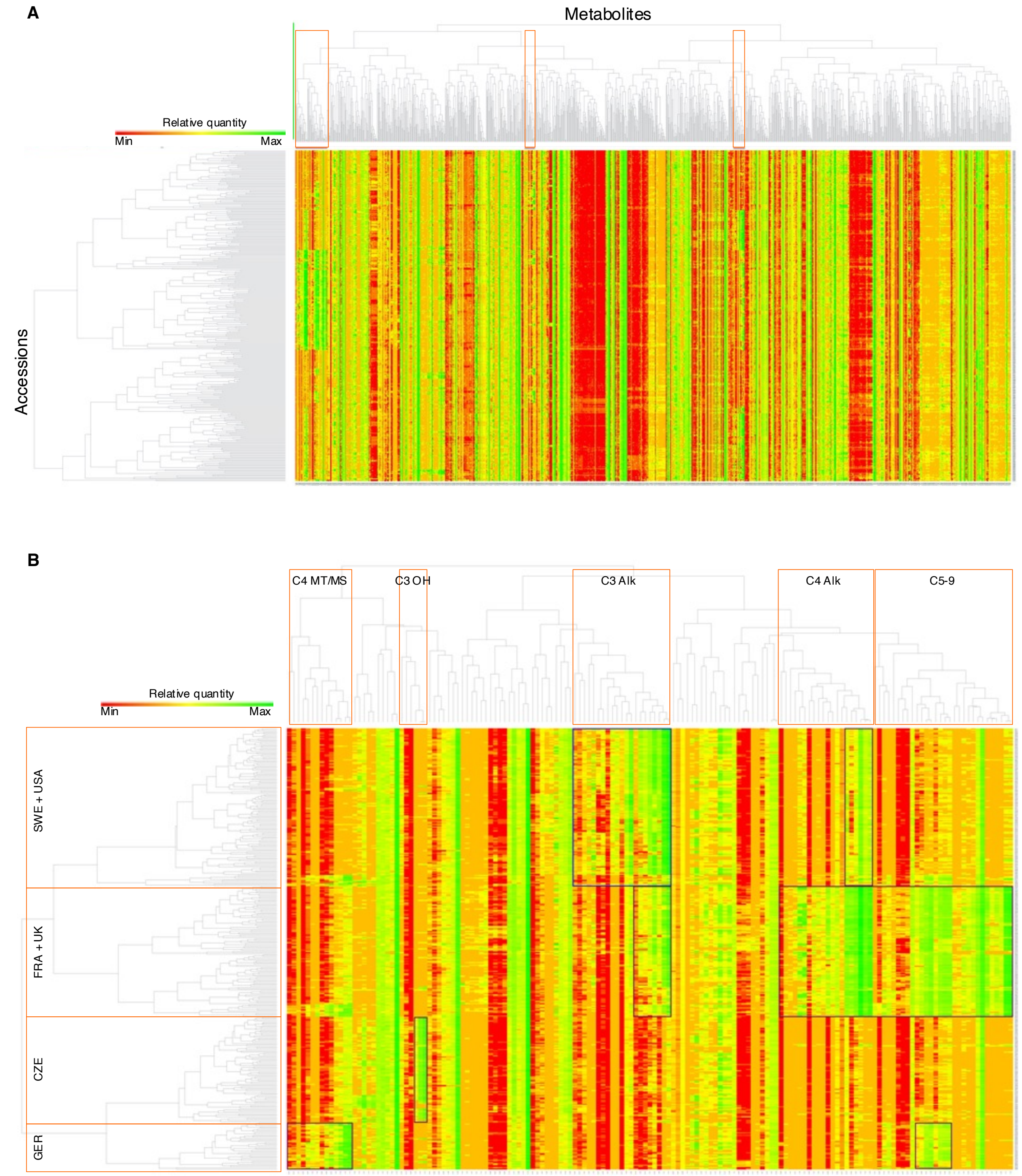


**Supplemental figure 8:** Two-dimensional hierarchical clustering of Arabidopsis accessions and their metabolic content.

(**A**) Pearson clustering of all detected and quantified volatile and non-volatile metabolites. Red boxes indicate glucosinolates and glucosinolate derivatives. (**B**) Pearson clustering of aliphatic glucosinolates only. Metabolites clustered in columns can be grouped in distinct glucosinolate classes, as indicated by the red boxes. Accessions clustered in rows could be grouped in four geographical regions of collection, as indicated by the red boxes.

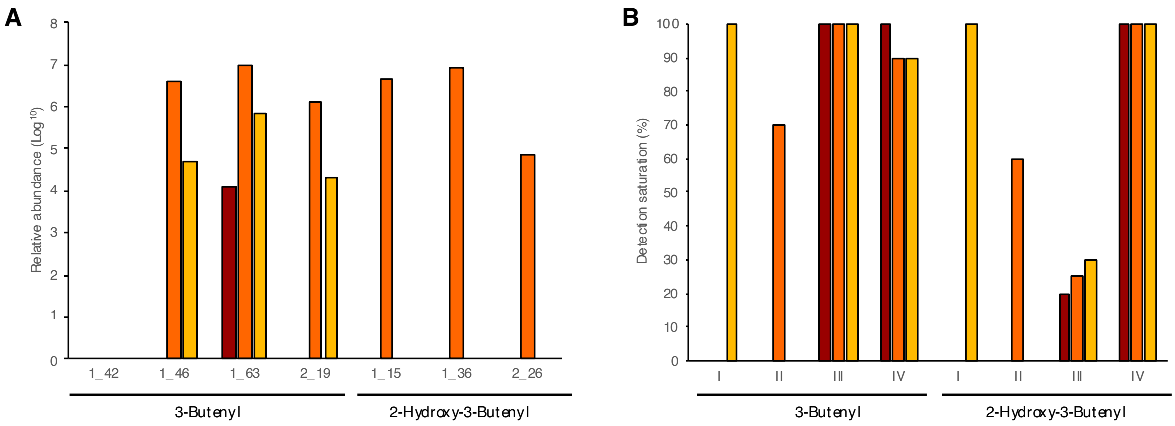


**Supplemental figure 9:** Detection of alkenyl glucosinolates in different genotypes of Arabidopsis.
(**A**) Relative abundance of masses putatively annotated as 3-butenyl (gluconapin) or 2-hydroxy-3-butenyl ((epi)progoitrin) detected in the Arabidopsis accessions Col-0, UKNW06-460 and Tamm-2 (ordered from dark to light bars), respectively. (**B**) Abundance of 3-butenyl and 2-hydroxy-3-butenyl, expressed as percentage of detection saturation, in doubled haploids of four haplotype classes derived from a cross between UKNW06-460 and Tamm-2. Three replicate pools of two plants each were tested for each haplotype, ordered from dark to light bars. The four haplotypes differ in functional alleles for the *GSOH* and *DMR6* loci (Haplotype I: *GSOH* silent, *DMR6* silent; Haplotype II: *GSOH* active, *DMR6* silent; Haplotype III: *GSOH* silent, *DMR6* active; Haplotype IV: *GSOH* active, *DMR6* active).

| SNP / Locus | 32846 ESP | 68088 GS-OH | 126970 AOP | 127038 AOP | 127050 AOP | 127067 AOP | 127105 AOP | 127113 AOP | 175356 MAM | 175357 MAM | 175362 MAM | 175369 MAM | 175386 MAM | 175661 TT16 | 175687 TT16 | 176930 DMR6 | 177048 DMR6 | 177054 DMR6 |
|---|---|---|---|---|---|---|---|---|---|---|---|---|---|---|---|---|---|---|
| 32846 ESP | | 0.17 | 0.10 | 0.02 | 0.05 | 0.05 | 0.05 | 0.05 | 0.17 | 0.19 | 0.20 | 0.15 | 0.14 | 0.00 | 0.00 | 0.06 | 0.14 | 0.15 |
| 68088 GS-OH | 0.17 | | 0.17 | 0.05 | 0.05 | 0.05 | 0.05 | 0.05 | 0.25 | 0.28 | 0.31 | 0.29 | 0.25 | 0.01 | 0.02 | 0.18 | 0.28 | 0.29 |
| 126970 AOP | 0.10 | 0.17 | | 0.23 | 0.12 | 0.11 | 0.12 | 0.14 | 0.23 | 0.24 | 0.21 | 0.22 | 0.19 | 0.00 | 0.00 | 0.14 | 0.17 | 0.11 |
| 127038 AOP | 0.02 | 0.05 | 0.23 | | 0.16 | 0.17 | 0.15 | 0.04 | 0.06 | 0.06 | 0.06 | 0.06 | 0.05 | 0.00 | 0.00 | 0.04 | 0.03 | 0.01 |
| 127050 AOP | 0.05 | 0.05 | 0.12 | 0.16 | | 0.95 | 0.92 | 0.62 | 0.13 | 0.13 | 0.11 | 0.12 | 0.08 | 0.01 | 0.01 | 0.03 | 0.04 | 0.06 |
| 127067 AOP | 0.05 | 0.05 | 0.11 | 0.17 | 0.95 | | 0.88 | 0.63 | 0.13 | 0.14 | 0.12 | 0.13 | 0.09 | 0.01 | 0.01 | 0.02 | 0.05 | 0.06 |
| 127105 AOP | 0.05 | 0.05 | 0.12 | 0.15 | 0.92 | 0.88 | | 0.59 | 0.13 | 0.14 | 0.12 | 0.12 | 0.09 | 0.02 | 0.02 | 0.02 | 0.04 | 0.06 |
| 127113 AOP | 0.05 | 0.05 | 0.14 | 0.04 | 0.62 | 0.63 | 0.59 | | 0.11 | 0.14 | 0.10 | 0.11 | 0.09 | 0.03 | 0.02 | 0.03 | 0.06 | 0.05 |
| 175356 MAM | 0.17 | 0.25 | 0.23 | 0.06 | 0.13 | 0.13 | 0.13 | 0.11 | | 0.86 | 0.78 | 0.72 | 0.63 | 0.12 | 0.12 | 0.10 | 0.19 | 0.20 |
| 175357 MAM | 0.19 | 0.28 | 0.24 | 0.06 | 0.13 | 0.14 | 0.14 | 0.14 | 0.86 | | 0.81 | 0.78 | 0.70 | 0.14 | 0.12 | 0.13 | 0.23 | 0.23 |
| 175362 MAM | 0.20 | 0.31 | 0.21 | 0.06 | 0.11 | 0.12 | 0.12 | 0.10 | 0.78 | 0.81 | | 0.78 | 0.72 | 0.14 | 0.14 | 0.11 | 0.21 | 0.20 |
| 175369 MAM | 0.15 | 0.29 | 0.22 | 0.06 | 0.12 | 0.13 | 0.12 | 0.11 | 0.72 | 0.78 | 0.78 | | 0.79 | 0.14 | 0.13 | 0.11 | 0.19 | 0.18 |
| 175386 MAM | 0.14 | 0.25 | 0.19 | 0.05 | 0.08 | 0.09 | 0.09 | 0.09 | 0.63 | 0.70 | 0.72 | 0.79 | | 0.16 | 0.15 | 0.09 | 0.17 | 0.14 |
| 175661 TT16 | 0.00 | 0.01 | 0.00 | 0.00 | 0.01 | 0.01 | 0.02 | 0.03 | 0.12 | 0.14 | 0.14 | 0.14 | 0.16 | | 0.83 | 0.01 | 0.00 | 0.01 |
| 175687 TT16 | 0.00 | 0.02 | 0.00 | 0.00 | 0.01 | 0.01 | 0.02 | 0.02 | 0.12 | 0.12 | 0.14 | 0.13 | 0.15 | 0.83 | | 0.01 | 0.01 | 0.01 |
| 176930 DMR6 | 0.06 | 0.18 | 0.14 | 0.04 | 0.03 | 0.02 | 0.02 | 0.03 | 0.10 | 0.13 | 0.11 | 0.11 | 0.09 | 0.01 | 0.01 | | 0.36 | 0.22 |
| 177024 DMR6 | 0.14 | 0.28 | 0.17 | 0.03 | 0.04 | 0.05 | 0.04 | 0.06 | 0.19 | 0.23 | 0.21 | 0.19 | 0.17 | 0.00 | 0.01 | 0.36 | | 0.66 |
| 177054 DMR6 | 0.15 | 0.29 | 0.11 | 0.01 | 0.06 | 0.06 | 0.06 | 0.05 | 0.20 | 0.23 | 0.20 | 0.18 | 0.14 | 0.01 | 0.01 | 0.22 | 0.66 | |

0 LD 1

**Supplemental figure 10:** Linkage disequilibrium between loci explaining variation in glucosinolate content. Each locus is represented by multiple SNPs, detected by BayesR WGR as the largest-effect SNP explaining variation in one or more glucosinolates.
